# Northward Expansion of *Amblyomma americanum* (Acari: Ixodidae) into Southern Michigan

**DOI:** 10.1101/2021.12.31.474651

**Authors:** Peter D. Fowler, S. Nguyentran, L. Quatroche, M.L. Porter, V. Kobbekaduwa, S. Tippin, Guy Miller, E. Dinh, E. Foster, J.I. Tsao

## Abstract

*Amblyomma americanum* (Linnaeus) (Acari: Ixodidae) (lone star tick) is an aggressive, generalist parasite that vectors numerous important human and animal pathogens. In recent decades its geographic range has expanded northwards from endemic regions in the southeastern and southcentral US. In 2019 five questing *A. americanum* comprising two life stages were detected at one site in Berrien County, in southwestern Michigan, satisfying one CDC criterium for an established population for the first time in the state. To better characterize the northern extent of emerging *A. americanum*, we conducted active surveillance (i.e., drag sampling) in summer 2020 throughout Michigan’s southern counties and detected one adult *A. americanum* from each of six widespread sites, including where they had been detected in 2019. A larger established population was identified at another site in Berrien County, which yielded 691 *A. americanum* comprising three life stages. Questing tick phenologies at this site were similar to that reported for other regions. Statewide surveillance in 2021 revealed no *A. americanum* outside of Berrien County, but establishment criteria were met again at the two sites where established populations were first detected respectively in 2019 and 2020. These observations may represent the initial successful invasion of *A. americanum* into Michigan. Data from passive (1999-2020) and active surveillance (2004-2021) efforts, including a domestic animal sentinel program (2015-2018), are reported to provide context for this nascent invasion. Continued active surveillance is needed to help inform the public, medical professionals, and public health officials of the health risks associated with this vector.

## Introduction

*Amblyomma americanum* (Linnaeus) (Acari: Ixodidae), also known as the lone star tick or turkey tick, is a generalist parasite known for its aggressive behavior (Stafford et al. 2018, Molaei et al. 2019). Aside from being a nuisance, it can bear negative health consequences. Heavy infestations of *A. americanum* on wildlife can cause morbidity and mortality (Bolte et al. 1970, Stafford et al. 2018). *Amblyomma americanum* is also known to vector a number of pathogens resposnsible for disease in both humans and domestic animals (Jaworski et al. 2017, Madison-Antenucci et al. 2020, Kennedy and Marshall 2021) and bites from this tick have also been linked with the recently discovered alpha-gal syndrome (AS), otherwise known as the tick-associated red meat allergy (Cabezas-Cruz et al. 2019, Crispell et al. 2019, de la Fuente et al. 2020, Sharma and Karim 2021), which, if serious, can result in anaphylaxis (Pattanaik et al. 2018).

Since the mid-1900s, the geographic range of *A. americanum* has been expanding northwards from endemic areas in the south central and southeastern US (Paddock and Yabsley 2007, Springer et al. 2014, Sonenshine 2018, Molaei et al. 2019). Management practices beginning in the early 20^th^ century to recover and increase forests and wildlife resulted in the growth and spread of its main host, the white-tailed deer (*Odocoileus virginianus*), and possibly the wild turkey (*Meleagris gallopavo*) (Paddock and Yabsley 2007). This increase in woodland habitats created more habitat not only for host species but also for *A. americanum* itself by providing leaf litter suitable to protect the tick from desiccation (Paddock and Yabsley 2007, Tsao et al. 2021). Additionally, the trend of warmer and shorter winters beginning in late 20^th^ century associated with anthropogenic climate change may be providing additional opportunities for this tick to become established in new areas farther north (Sonenshine 2018). Models suggest that the northward invasion of this tick is likely to continue in the coming decades (Springer et al. 2015, Ludwig et al. 2016, Raghavan et al. 2019, Sagurova et al. 2019). Detecting, further characterizing, and refining predictions about the northward expansion can help prepare the public and medical professionals for the emergence of diseases associated with this vector.

Recently, breeding populations of *A. americanum* have been documented in southern New England (Stafford et al. 2018, Telford et al. 2019) and in states surrounding the Great Lakes, established populations of have been spreading northwards in Illinois (Gilliam et al. 2020, Lyons et al. 2021), Indiana (Wojan et al. 2021), and Ohio (Fitak et al. 2014). Although there have been increases in *A. americanum* detected by the public in Wisconsin (Christenson et al. 2017) and southern Ontario (Nelder et al. 2019), established populations have not yet been declared in those areas (at the time of manuscript submission). As of July 2019, the Minnesota Department of Public Health has not reported any established populations of *A. americanum* although they have been reported by residents since 1998 (https://www.health.state.mn.us/diseases/tickborne/longstarstickreportsmap.pdf, accessed 10/6/21), and established populations have been identified in nearby counties to the west in South Dakota (Black et al. 2021) and to the south in Iowa (Springer et al. 2014).

Although *A. americanum* in Michigan have been documented by passive surveillance since the mid-1980s (Walker et al. 1998, MDHHS 2021a) there is a lack of evidence supporting the existence of any established breeding populations (Walker et al. 1998). Here we present such data for one county in Michigan. To better understand the context of this invasion, we also summarize detections of *A. americanum* reported in historical passive and active surveillance efforts dating back to 2004 designed to detect *Ixodes scapularis* (Say) because there is a high degree of overlap between *A. americanum* and *I. scapularis* in habitat use and temporal activity (Lindsay et al. 1998, Kollars et al. 2000, Guerra et al. 2002, Allan 2009, Centers for Disease Control and Prevention 2020). We further include data from active surveillance also focused on detecting *I. scapularis* carried out by the Berrien County Health Department (BCHD) from 2019-2021. Finally, we present data from a broad statewide surveillance study carried out in 2010 in order to define the range of the then emerging *I. scapularis*.

## Methods

### Passive surveillance

Records of resident-submitted ticks to the Michigan Department of Agriculture began in 1999. In 2015 the Michigan Department of Health and Human Services (MDHHS) took over the program, which continues to the present. Data were retrieved from the public database on the MDHHS Michigan Environmental Public Health Tracking website (MDHHS 2021a). Ticks are reported by the submitter’s most likely county of exposure or county of residence when the county of exposure is uncertain. Submitted ticks were identified to species using dichotomous morphological keys (Brinton et al. 1965, Sonenshine 1979, Keirans and Litwak 1989, Durden and Keirans 1996, Faulkner and Reinhard 2014, Dubie et al. 2017, Egizi et al. 2019). Although the program was in operation in 2014, data were not available for this analysis.

### Active surveillance by drag sampling

#### Site selection

From 2004-2016, much of the sampling by Michigan State University (MSU) was conducted to track the invasion of *I. scapularis* mainly in the western Lower Peninsula (Hamer et al. 2010), but this was expanded to southeastern Michigan beginning in 2015. In 2010 in order to better define the range of *I. scapularis* surveillance was broadened to cover 21 counties across Michigan’s lower penninsula. Beginning in 2017, surveillance became more widespread across the state in order to sample counties that previously had been sampled less frequently or not at all. In 2019 BCHD began conducting surveillance for questing *I. scapularis.* Although the specific monitoring objectives varied from year to year, site selection and sampling protocols were similar between MSU and BCHD. Public land (e.g., city, county and state parks, state forests, state wildlife game areas, and national forests) with suitable habitat for *I. scapularis* were selected. Much like *I. scapularis, A. americanum* is frequently found in deciduous forests containing leaf litter that helps protect ticks from desiccation (Hair and Howell 1970, Childs and Paddock 2003, Allan 2009, Kensinger and Allan 2011). In 2020, the number of sites per county sampled by MSU was increased relative to that in prior years in order to increase the likelihood of detecting *A. americanum* in southern Michigan: fifty-five primary sites were selected after consulting local public health departments, local parks departments and other regional authorities to identify sites with suitable habitat and public access. Sites with insufficient area to sample (less than 1000 meters) or unsuitable habitat (little to no deciduous forest, or lack of a managed trail) upon first visits were not repeated.

#### Surveillance methods and sampling frequency

Active surveillance by drag sampling (Falco and Fish 1992) aimed at detecting the spread of *I. scapularis* has been ongoing in Michigan since 2002 (Foster 2004). Based on CDC guidelines for the active surveillance of questing prostriate (Eisen et al. 2019) and metastriate ticks (Centers for Disease Control and Prevention 2020), a minimum of 1,000 meters were sampled per visit per site by dragging a 1-m^2^ white corduroy cloth along trail edges in contact with understory and leaf litter where ticks were likely questing. In larger parks, two transects with a minimum of 800 meters each were sampled in different areas of the park. Sampling was performed on rain-free days throughout the day unless relative humidity dropped below 40%. Collected ticks were placed in 70-90% ethanol and brought back to the lab for species identification using dichotomous morphological keys (Brinton et al. 1965, Sonenshine 1979, Keirans and Litwak 1989, Durden and Keirans 1996, Faulkner and Reinhard 2014, Dubie et al. 2017, Egizi et al. 2019).

In 2004-2019 and 2021, because the main objective of our surveillance was to document the distribution of *I. scapularis* and Lyme disease risk, the bulk of the sampling effort generally occurred from May - July to target the nymphal stage of this tick. This time frame also encompasses part of adult and much of the nymphal host-seeking period of *A. americanum*, according to reported phenologies in other states (Sonenshine 1979, Schulze et al. 1986, Kollars et al. 2000, Telford et al. 2019). Sites were sampled at least once during these years. In 2020, COVID-19 restrictions delayed sampling efforts by MSU, but sites with appropriate habitat and area were sampled a minimum of three times from June 8th - July 23rd, 2020. Sites where *A. americanum* was detected during the first three visits were sampled an additional time (i.e., four times total) to attempt to detect multiple life stages or multiple specimens. Figures were generated in ArcGIS Pro v. 2.8.3 and compiled in Adobe Illustrator v. 24.1.2.

### Active surveillance by sampling companion canines

From 2015-2018 veterinary clinics and/or shelters throughout Michigan were recruited to survey companion canines for ticks during routine health checks. Veterinary workers were asked to inspect thoroughly a minimum of 60 patients (3 per day) over a period of at least 30 days when adult *I. scapularis* ticks would be active in Michigan, targeting April - June. *Amblyomma americanum* frequently parasitizes companion canines (Sonenshine 1979, Saleh et al. 2019) and this timeframe overlaps the activity period of the adult *A. americanum* as well as part of that of the nymph. Only canines who had not traveled more than 30 miles from their residence during the 10 days prior to their health visit were included in the study. Clinics were instructed to report data on all canines inspected, including those who did not present with ticks. All ticks were stored in 70% ethanol and submitted to Michigan State University for identification as described above. The activities herein are considered standard veterinary care and involve samples collected using non-invasive techniques; an approved exemption (4/20/15) from filing an animal use form is on file with the MSU Institutional Animal Care and Use Committee.

### *Characterization of the phenology of* A. americanum

Because both adults and nymphs were detected at one site, Grand Mere State Park (GMSP), early in the surveillance period in 2020, weekly sampling beyond the surveillance window was performed there to detect larvae as well as to describe the host-seeking phenology of *A. americanum*. Sampling in the fall ceased after two consecutive visits failed to yield any *A. americanum*. Host-seeking phenologies for each tick life stage was characterized by determining the density of ticks per site visit in ticks per m^2^ and displaying the percentage of the total tick density for the season for each respective visit.

## Results

### Passive Surveillance

Figure 1 shows the numbers of ticks submitted by county to the Michigan Department of Agriculture from 1999-2014 and MDHHS from 2015-2020 excluding 2014 for which data were not available. Figure 2 shows the total number of ticks submitted from 2009 to 2020; *A. americanum* submissions accounted for 1.8% (0.6 - 3.0% range) of all ticks submitted each year. Submissions were variable in space and time with no apparent trends especially given the spatial distribution of Michigan residents.

**Figure 1.**
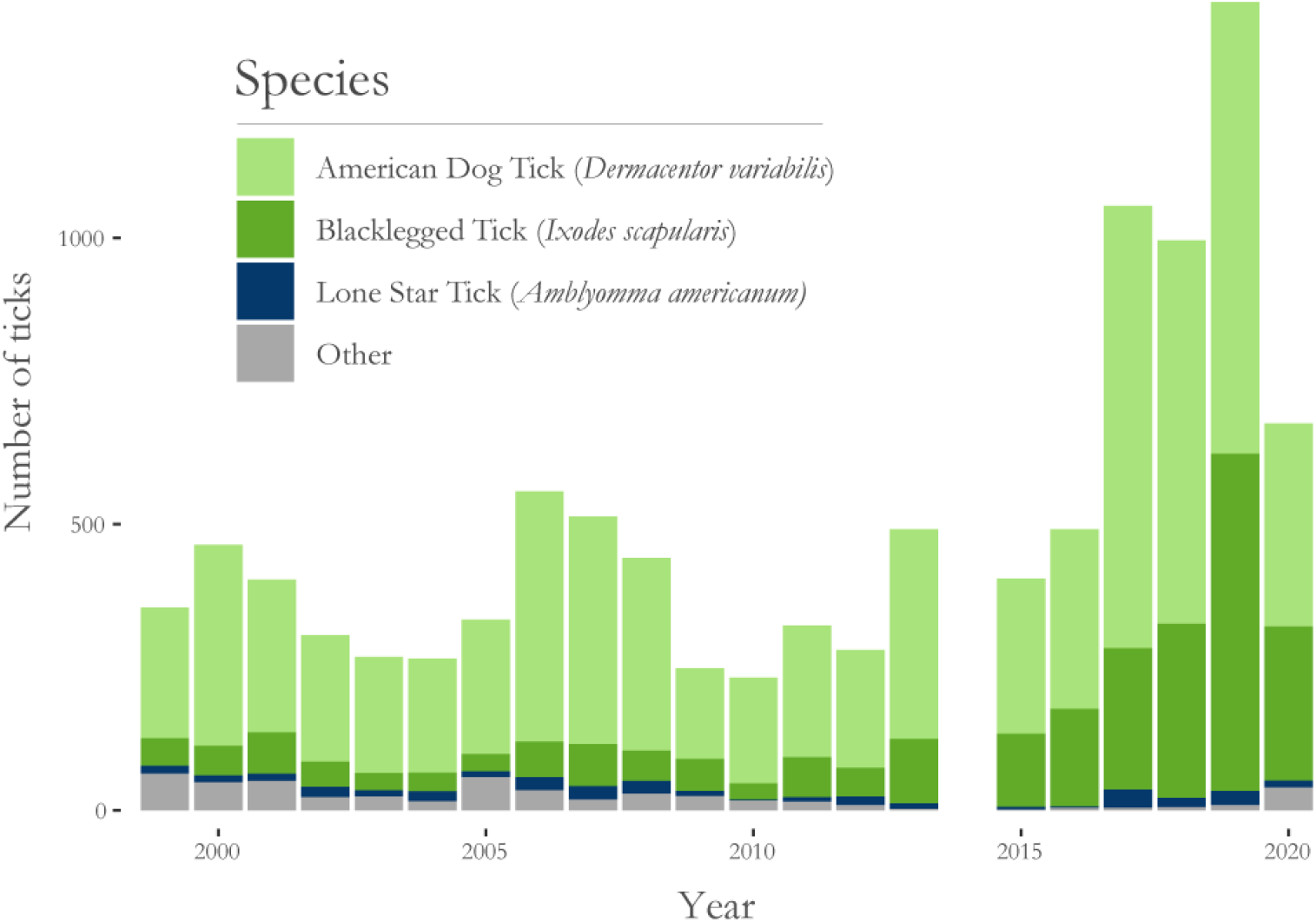
Numbers of *Dermacentor variabilis, Ixodes scapularis, Amblyomma americanum*, and other species submitted annually to the Michigan Department of Agriculture (1999 - 2013) and Michigan Department of Health and Human Services (2015 - 2020). On average, 3.2% ±1.7% SD of all ticks submitted annually were *A. americanum*. Data source: MiTracking-Michigan Environmental Public Health Tracking; no data are available for 2014.

**Figure 2.**
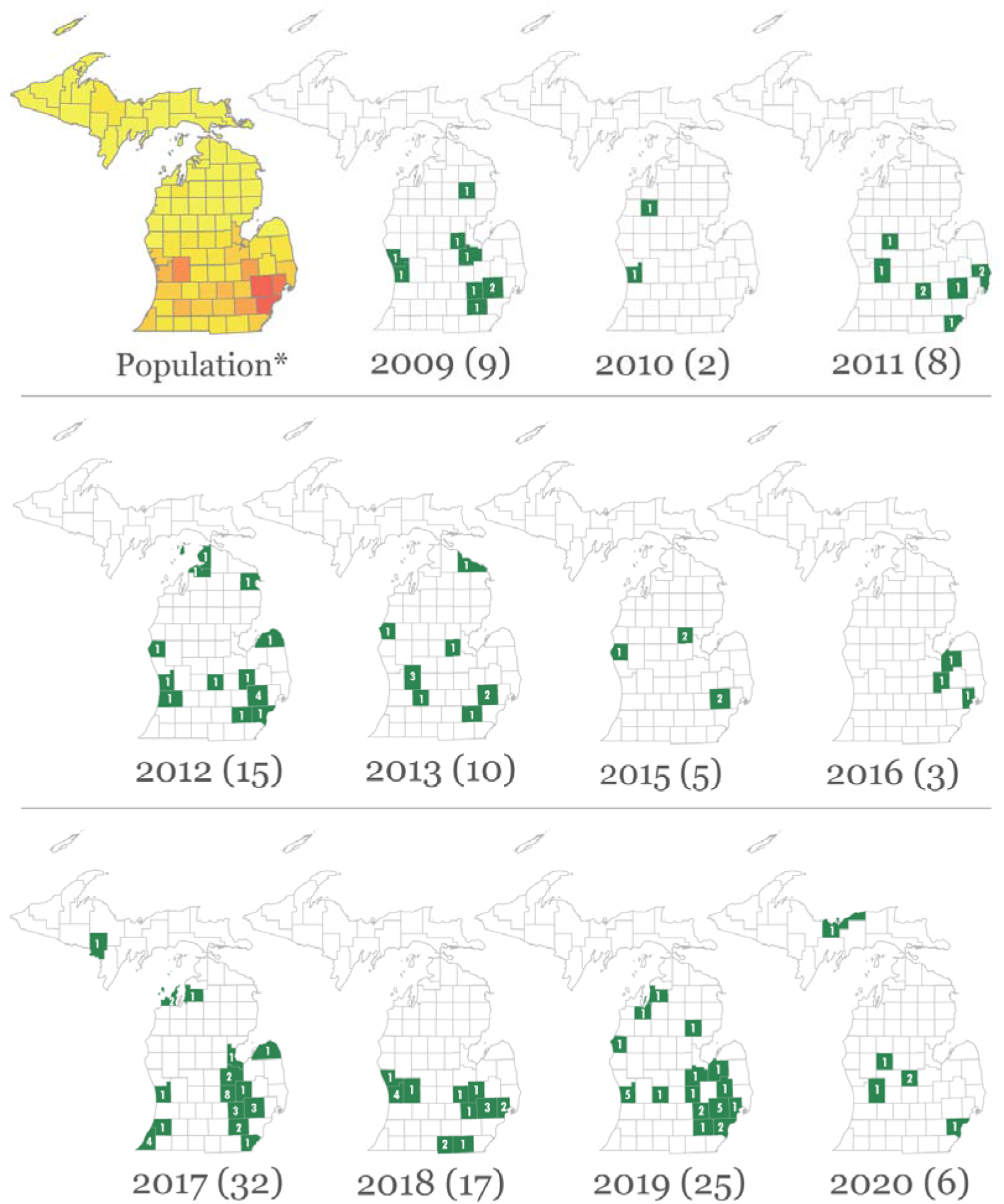
*Amblyomma americanum* submitted annually to the Michigan Department Agriculture (2009-2013) and the Michigan Department of Health and Human Services (MDHHS) from 2015-2020. Data source: MiTracking-Michigan Environmental Public Health Tracking. Numbers in each county represent the number of community-submitted *A. americanum* by county of residence. No data were available for 2014. * Human population heatmap for each county is given for reference based on 2020 census date (collated by Esri 2021) with lighter yellow being lower population and darker orange being higher population.

### Active Surveillance

#### Drag sampling (2004-2021)

Between 2004 and 2016, 628,140 m^2^ were sampled from 35 counties located in the Lower Peninsula (Allegan, Barry, Bay, Benzie, Berrien, Calhoun, Cass, Charlesvoix, Clinton, Ingham, Ionia, Iosco, Iron, Isabella, Jackson, Kalamazoo, Kent, Lapeer, Leelanau, Livingston, Manistee, Mason, Montcalm, Muskegon, Oakland, Oceana, Osceola, Ottawa, Roscommon, Saginaw, Sanilac, St. Clair, Tuscola, Van Buren, Wayne) (Hamer et al. 2010 (2004-2008); Tsao, unpublished data (2009-2016)). Four *A. americanum* were recovered, all from southwestern Michigan: one adult female in Barry County (2004), one adult female from Allegan County (2005), one adult female in Van Buren County (2005), and one nymph in Barry County (2007) (Table 1).

**Table 1.** Questing *Amblyomma americanum* detected by active surveillance (drag sampling) in Michigan from 2004 - 2021 by life stage and county. Sampling in 2004-2016 was focused on the western side of the state (with a few exceptions - 2010 was broadened to better identify the range of *I. scapularis*). Beginning in 2017 active surveillance was carried out statewide except in 2020 when surveillance efforts focused on southern counties. See text for details. Only years where *A. americanum* were detected are included.

| Year | County | No. of *Amblyomma americanum* detected |  |  |  |
| --- | --- | --- | --- | --- | --- |
|  |  | Adult Male | Adult Female | Nymph | Larvae |
| 2004 | Barry | - | 1 | - | - |
| 2005 | Allegan | - | 1 | - | - |
|  | Van Buren | - | 1 | - | - |
| 2007 | Barry | - | - | 1 | - |
| 2017 | Van Buren | 1 | - | - | - |
| 2019 | Berrien | 1 | - | 4 | - |
|  | Muskegon | - | 1 | - | - |
| 2020 | Berrien | 1 | 3 | 32 | 409 |
|  | Branch | 1 | - | - | - |
|  | Huron | 1 | - | - | - |
|  | Ingham | 1 | - | - | - |
| 2021 | Berrien | 5 | 1 | 45 | 50 |
| Total |  | 11 | 8 | 82 | 459 |

Figure 3 shows the distribution of sampling effort across Michigan from 2017-2021 including the efforts by BCHD (2019-2021, Berrien County). Ingham County has a site marked by a yellow star which was sampled at regular intervals throughout the season in order to monitor regional activity of the target species, normally *I. scapularis*; only the first three visits to this site are included in the effort calculations as these efforts do not represent surveillance across the county and would skew effort reported in the figure (Figure 3). From a total of 212,989 m^2^ sampled in 2017, only one adult male *A. americanum* was recovered (Barry County). From 467,688 m^2^ sampled in 2018, no *A. americanum* was detected. From 511,330 m^2^ sampled in 2019, five *A. americanum* of two life stages (1 adult male and 4 nymphs) were recovered at a single site in Berrien County, fulfilling one of the CDC criteria for an established population (Centers for Disease Control and Prevention 2020). BCHD sampled 12,000 m^2^, that same year (2019) but no other *A. americanum* were detected. In 2020, a total of 265,745 m^2^ were sampled at 55 sites across 24 counties in southern Michigan by MSU between June 8^th^ and July 23^rd^. A total of 2,918 ticks were collected (Table 2) comprised of *I. scapularis* (90.3%), *Dermacentor variabilis* (Say) (7.8%), *A. americanum* (1.2%), *I. dentatus* (Marx) (0.01%), and *Haemaphysalis leporispalustris* (Packard) (0.1%). Of the 55 sites sampled, *A. americanum* was found at 7 sites in five counties across southern Michigan (from west to east: Berrien, Branch, Cass, Ingham, and Huron). Only one site, located in Berrien County, yielded more than one *A. americanum* and multiple life stages. An additional I 25,000 m^2^ were sampled from May 11^th^ to July 30^th^ in 2020 by the Berrien County Health Department, which revealed a single *A. americanum* from two sites. In 2021, statewide surveillance by MSU for *I. scapularis* resumed; 373,378 m^2^ were sampled and an additional 38,000 m^2^ were sampled by BCHD. *Amblyomma americanum* was only detected in Berrien County, where multiple ticks were detected at two sites, Grand Mere State Park (GMSP) and Warren Dunes State Park (WDSP).

**Figure 3.**
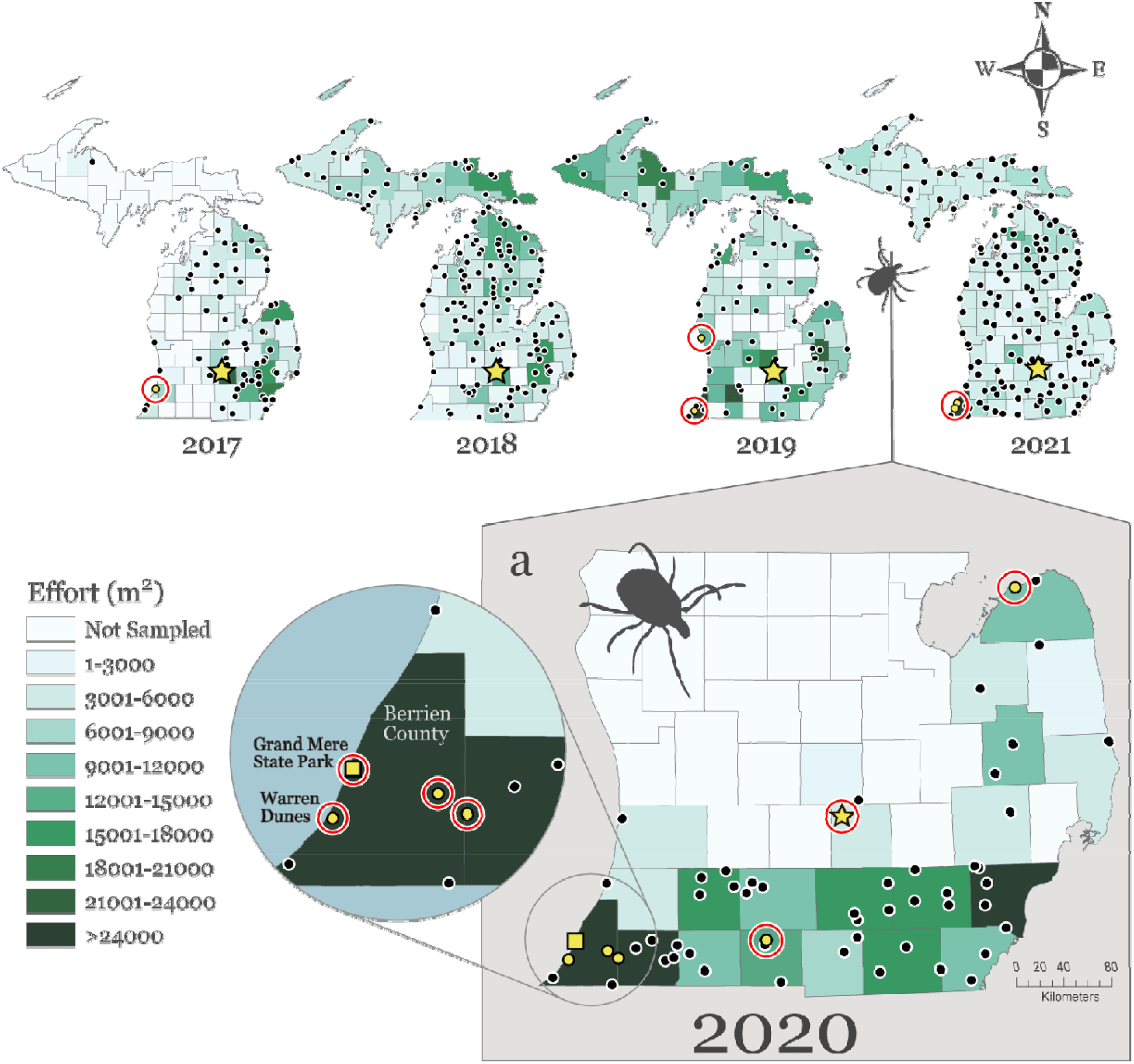
Drag sampling effort by county (m^2^) from 2017-2021. Black dots mark field sampling sites. Yellow dots with red circles mark sites where *A. americanum* were found. A) Sampling effort for 2017, 2018, 2019, 2021; B) Sampling effort for 2020 from June 8 - July 23 (See Table 1). The yellow square marks the established population of *A. americanum* at Grand Mere State Park (detected in this study). Yellow star marks a site sampled regularly as a reference for *I. scapularis* in order to ensure they are active regionally when sampling other sites. Only the first three visits for this site (i.e., during the nymphal host-seeking period) were included in the effort calculations for each year so as not to skew overall county effort.

**Table 2.** Total numbers of each tick species and life stage collected by drag sampling by county from June 8^th^ - July 23^rd^, 2020 in southern Michigan. AM = Adult Male; AF = Adult Female; N = Nymph; L = Larva; *H. lep. = Haemaphysalis leporispalustris; I. dentatus = Ixodes dentatus; I. cookei = Ixodes cookei; D. variabilis = Dermacentor variabilis; Ixodes* spp. = Unidentifiable *Ixodes* species due to damage to specimen.

|  |  | Species |  |  |  |  |  |  |  |  |  |  |
| --- | --- | --- | --- | --- | --- | --- | --- | --- | --- | --- | --- | --- |
|  |  | *Ixodes scapularis* |  |  |  | *Dermacentor variabilis* |  |  | *Amblyomma americanum* |  |  | Other |
| County | Effort (m²) | AM | AF | N | L | AM | AF | N | AM | AF | N |  |
| Allegan | 5,381 | 1 | 7 | 8 | - | 7 | 8 | - | - | - | - |  |
| Barry | 5,625 | - | - | 16 | 23 | - | - | - | - | - | - |  |
| Berrien | 42,039 | 26 | 26 | 278 | 309 | 22 | 32 | 4 | 3 | 3 | 26 | *I. dentatus* 1L, 3N |
| Branch | 12,529 | - | - | 2 | - | 1 | 1 | - | 1 | - | - |  |
| Calhoun | 10,855 | 5 | 3 | 52 | 1 | 3 | 2 | - | - | - | - | *I. dentatus* 2N |
| Cass | 25,161 | - | 3 | 128 | 835 | 1 | 8 | - | - | 1 | - | *I. dentatus* 1N |
| Clinton | 1,870 | 1 | - | 4 | 1 | - | - | - | - | - | - |  |
| Hillsdale | 8,060 | - | - | 8 | 31 | 4 | 4 | - | - | - | - |  |
| Huron | 10,277 | 1 | 4 | 3 | - | - | - | - | 1 | - | - | *H. lep.* 1N |
| Ingham | 4,700 | 8 | 7 | 250 | 82 | - | - | - | 1 | - | - |  |
| Jackson | 16,513 | 3 | - | 36 | 9 | 1 | 3 | - | - | - | - |  |
| Kalamazoo | 12,850 | 2 | 3 | 184 | 31 | 1 | 3 | - | - | - | - | *I. dentatus* 3N |
| Lapeer | 10,999 | - | - | 3 | 2 | - | 1 | - | - | - | - | *I. dentatus* 1N |
| Lenawee | 20,487 | - | - | 3 | - | 12 | 20 | - | - | - | - | *I. cookei* 1AF |
| Monroe | 11,336 | - | - | - | - | 10 | 7 | - | - | - | - | *I. dentatus* 1N |
| Oakland | 5,678 | 1 | 1 | 3 | - | 1 | - | - | - | - | - |  |
| Sanilac | 1,560 |  |  |  |  |  |  |  |  |  |  |  |
| St. Clair | 4,986 | - | 1 | 28 | 3 | - | - | - | - | - | - |  |
| St. Joseph | 9,300 | - | - | 63 | - | 3 | 7 | - | - | - | - | *I. dentatus* 1N |
| Tuscola | 4,296 | - | - | 5 | - | - | - | - | - | - | - | *Ixodes spp.* 1L |
| Van Buren | 5,732 | 5 | 3 | 52 | 6 | 21 | 30 | - | - | - | - | *H. lep.* 1N |
| Washtenaw | 10,230 | 5 | 3 | 25 | 5 | 1 | 1 | - | - | - | - | *I. dentatus* 1N |
| Wayne | 26,806 | 1 | - | 21 | 6 | 3 | 5 | - | - | - | - | *D. variabilis* 1L |
| Total | 265,745 | 59 | 61 | 1172 | 1344 | 91 | 132 | 4 | 6 | 4 | 26 | 19 |

#### Berrien County Active Surveillance

Table 3 summarizes the total sampling effort in Berrien County and numbers of *A. americanum* detected from 2019 - 2021 at all the sites that were sampled. Four parks were sampled all three years; two other parks were sampled only once. From the four parks sampled each year, *A. americanum* was detected at three sites (Table 3). At one site, Warren Dunes State Park (WDSP), the status of *A. americanum* was established in 2019 and was confirmed in 2021. Surprisingly only one *A. americanum* was found at this site in 2020 despite similar effort. At a second site, Grand Mere State Park (GMSP), *A. americanum* was not detected in 2019, but then multiple stages, including larvae, were detected in both 2020 and 2021, suggesting the presence of an established and breeding population. At the third site, Love Creek Nature Center, *A. americanum* were detected in only one year, in which two adults were detected thus not reaching the threshold criteria for an established population. Total sampling effort per site for all three years ranged from 26,003 m^2^ to 57,165 m^2^, and the total distance sampled at WDSP and GMSP was ~1.5-2 times more than the other two sites, where no *A. americanum* were detected or reported.

**Table 3.**
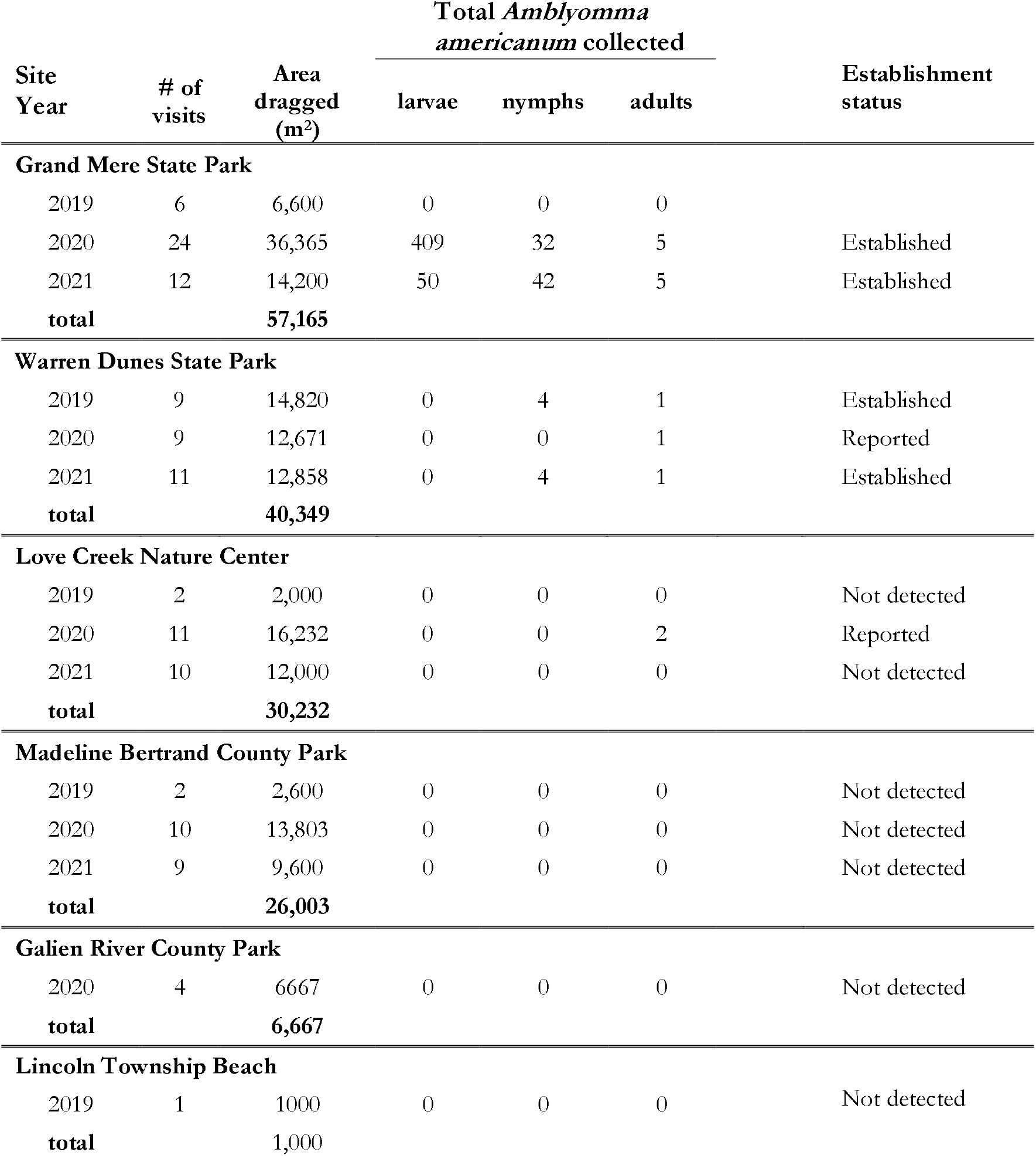
Active surveillance effort for Berrien County from 2019-2021and status level for *Amblyomma americanum* detections (per site per year) as defined by the CDC as more than 6 ticks or more than one life stage found in a given calendar year (Springer et al. 2014). Total effort for each site is reported for each year and includes sampling performed by both Michigan State University and the Berrien County Health Department. Two sites were sampled only for one year. Total numbers of *A. americanum* are reported for each site for each year. (Note: the lone star status of Berrien County as a whole would be considered established as of 2019, given the findings at Warren Dunes State Park.)

#### Sampling companion canines (2015-2018)

Study periods varied over the four years, but all included the periods when adult and nymphal *A. americanum* would be expected to be active (Table 4). At least 300 animals representing 16 - 69 counties per year were examined for a total of 4,097 animals examined. No *A. americanum* were detected in three of the four study years. In 2016, when 2,510 individuals were examined, a single adult female *A. americanum* was detected from a dog residing in Muskegon County.

**Table 4.** On-host *Amblyomma americanum* detected by active surveillance of companion canines with no travel history in Michigan from 2015 - 2018 by participating veterinary clinics or shelters.

|  | Year |  |  |  |
| --- | --- | --- | --- | --- |
|  | 2015 | 2016 | 2017 | 2018 |
| Dates animals were examined | 5/1 – 6/5 | 2/10 – 6/3 | 4/3 – 7/26 | 5/13 - 7/27 |
| No. clinics/shelters submitting data | 30 | 48 | 26 | 8 |
| No. of animals examined | 391 | 2519 | 887 | 300 |
| No. of counties represented | 31 | 69 | 42 | 16 |
| No. of animals infested with *A. americanum* | 0 | 1<sup>a</sup> | 0 | 0 |

<sup>a</sup>One adult female *A. americanum* was detected from an animal residing in Muskegon County.

### *Characterization of the phenology of* A. americanum *at GMSP*

A total of 448 *A. americanum* were sampled at GMSP between June 11^th^ and October 12^th^ 2020, including 409 larvae, 32 nymphs, 3 adult males, and 2 adult females. Seasonal activity for *A. americanum* and the now endemic *I. scapularis* (for comparison) are presented in Figure 4. *Amblyomma americanum* adults were detected only in June. Nymphs were active from June to September but were most active in June. Larvae were first detected in August but peaked in September and were not detected in October. *Ixodes scapularis* adults were detected in June, late September, and October, with greatest numbers in October. Nymphal *I*. *scapularis* were detected from June to September and peaked in June. Larval *I. scapularis* were detected throughout the study period, with a first peak in July followed by a second peak in September.

**Figure 4.**
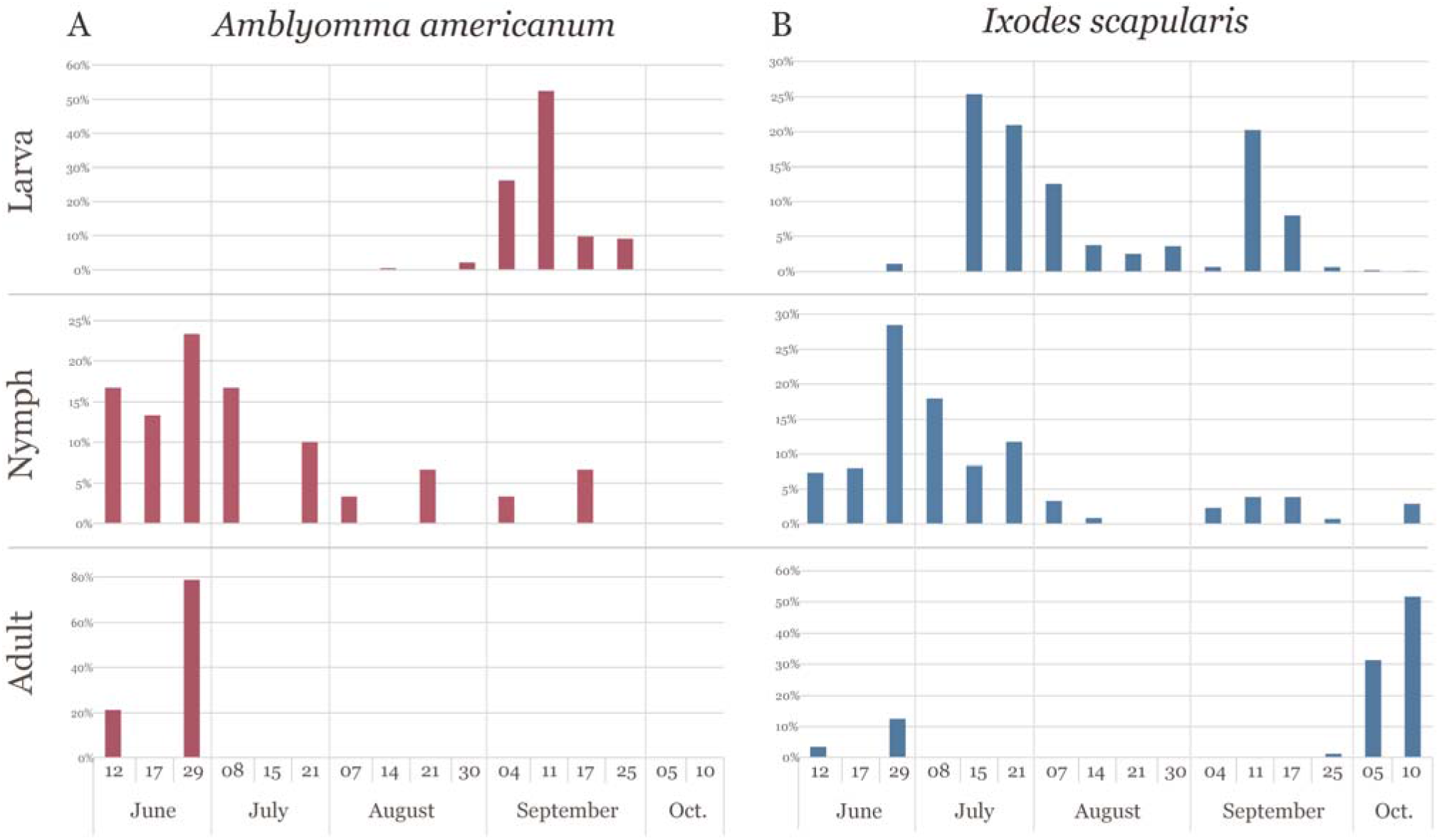
Phenology of each questing life stage of A) *Amblyomma americanum* and B) *Ixodes scapularis* at Grand Mere State Park (Berrien County) from 6/11/20 - 10/10/20 (ended after 2 consecutive visits in the fall where no *A. americanum* were detected). See the yellow square on the map in Figure 2 for the location of the field site.

## Discussion

Since the mid-1980s, detections of *A. americanum* in Michigan have been recorded in passive surveys (Walker et al. 1998, MDHHS 2021a); however, until recently, active surveillance efforts only rarely detected *A. americanum.* Applying the CDC’s operational definition for established tick populations (Springer et al. 2014), we report the first records of established populations of *A. americanum* in Michigan, and according to CDC convention, the lone star tick status of Berrien County should be considered established since 2019. To provide evidence that this invasion is recent and geographically limited we collated the results of prior surveillance efforts in Michigan comprising passive surveillance for ticks in general and active surveillance for blacklegged ticks specifically. Weekly surveillance at one site where a breeding population of *A. americanum* was detected allowed us to characterize the seasonal activity of this newly establishing tick in Michigan, which appears to be similar to that in other areas where lone star ticks are established.

### Previous active surveillance in Michigan (2004-2019)

Between 2004 and 2018, 841,129 m^2^ were sampled throughout the state and only four adult and one nymphal *A. americanum* were detected despite sampling in suitable habitats during phenologically appropriate times of the year (Hamer et al. 2010; Tsao, unpublished data). While these efforts may have missed some cryptic populations, especially in less sampled regions (e.g, the Upper Peninsula), the findings suggest that there were no widespread established populations of *A. americanum* within Michigan during this timeframe. Furthermore, examination of > 4,000 companion canines over four years (2015-2018) across Michigan resulted in the detection of one *A. americanum* tick attached to one dog, who resided in a county in the Lower Peninsula along Lake Michigan.

In 2019, drag sampling revealed five *A. americanum* of two life stages (one adult male and four nymphs) all from Warren Dunes State Park (WDSP). We took immediate notice of this finding because detecting five ticks at one site in one summer starkly contrasts with detecting a total of five ticks from three counties over fifteen years. We also took notice because prior detections mainly were comprised of single adult ticks, whereas at WDSP in 2019, 4 of 5 ticks were nymphs. For nymphs to be detected in an area, they may have derived from locally hatched larvae that fed on local hosts, thereby representing a locally reproducing population. On the other hand, they may have derived from larvae that had hatched elsewhere and then were introduced into the area by dispersing hosts. For example, animals, such as deer, coyote, or birds, dispersing from northwest Indiana or other areas in southwest Berrien County with undetected lone star tick populations may have introduced larvae. In either of these cases, introduction of larvae would be from within the same local region, suggesting the successful invasion lone star ticks in Michigan. Considering that lone star tick larvae generally host-seek in late summer (Telford et al. 2019), it is unlikely that migratory songbirds would have introduced larvae from the endemic regions in southeastern or south central U.S. or even populations in southern Indiana or Illinois, as fall migrating songbirds are traveling in the opposite direction. Although it is possible for nymphs to be dispersed northwards by spring migrating songbirds, as spring migration overlaps with the early part of the nymphal host-seeking period, multiple successful events as well as surviving the local abiotic conditions would need to follow, for these engorged nymphs to result in local larval ticks (e.g., molting, finding an adult host, finding a mate, ovipositing, and larval hatch). A high volume of migrating birds landing and/or stopping over in an area may elevate the chances of such introductions. However, introductions of dispersing larvae, nymphs or adults from nearby established populations might be more likely.

### *Targeted surveillance for* A. americanum *in Michigan in 2020 and statewide surveillance in 2021*

Given the finding of multiple life stages of *A. americanum* at WDSP in southwestern Michigan, and given the presence of established *A. americanum* populations in northwestern Indiana counties that border southwestern Michigan, (L. Green, Indiana Department of Public Health (IDOH) pers. comm.) surveillance conducted by MSU in 2020 focused on southern counties. *Amblyomma americanum* were detected at a greater number of sites and counties in 2020 than in any previous year, and this may have been due in part to greater sampling effort per county compared with that in previous years. First, although sampling effort per site was similar in 2020 compared to 2019 (approximately 3-4 visits/site), it was generally greater compared to that in prior years. Second, in 2020 we attempted to increase the concentration of sites from ~1-2 sites per county to 3-5 sites per county. Even with this increased coverage of southern Michigan, only one county (Berrien) yielded more than one site with a *A. americanum* tick detection, and only one site, Grand Mere State Park (GMSP) in Berrien County, revealed more than one life stage. It is not surprising that sampling a greater density of sites may result in a greater number of detections of *A. americanum*. Although the sample size is small (n = 5), interestingly, the detections outside of Berrien County were all adults, which may represent adventitious nymphs dispersed by from endemic areas in the south central and southeastern U.S. by spring migrating songbirds (as discussed in Stafford et al. 2018). At one of the sites outside Berrien County where an adult *A. americanum* was detected in 2020, ~ bi-weekly sampling May - October in 2019 and 2021 did not result in any detections of *A. americanum.*

### *A. americanum* tick emergence in Berrien County

In the early 2000s Berrien County was sampled frequently because it is also where established populations of *I. scapularis* were first detected in the Lower Peninsula (Foster et al. 2004). Because of the limited resources and the need to monitor other areas where the establishment status of *I. scapularis* was less known, Berrien County was not sampled as intensively (i.e., number of sites and frequency of visits per year) by MSU after the early 2000s. Beginning in 2019, sampling effort in Berrien County increased in large part due to the initiation of surveillance by BCHD as part of a MDHHS-funded program to increase surveillance efforts for invasive *I. scapularis* and pathogens in Michigan. MSU efforts also increased in 2020 following the discovery of multiple life stages of *A. americanum* at WDSP in 2019. Subsequently, multiple life stages of *A. americanum* were detected in two years for two sites, WDSP and GMSP.

Although *A. americanum* were collected in each year (2019-2021) at WDSP, the tick collections only met one of the CDC criteria for established populations (detecting at least two life stages in one calendar year) in 2019 and 2021. The numbers of ticks collected each year were low, never achieving the other criterium of detecting at least 6 ticks. In contrast, data from GMSP suggest that *A. americanum* either were not yet established at GMSP in 2019 or were present at an abundance below our threshold of detection, but then appear to be steadily increasing in 2020 and 2021. Taken together, established populations, according to the CDC criteria and applied at the site level, have been detected in Berrien County for three consecutive years.

No established populations of *A. americanum* were detected outside WDSP and GMSP, suggesting that either further spread has not yet occurred, or that the level of sampling effort was inadequate to detect low density emerging populations. Future research may investigate differences in abiotic and biotic factors among sites to understand variation in initial establishment success of *A. americanum.* For instance, both WDSP and GMSP are located along Lake Michigan, whereas other sites are found further inland. Proximity to Indiana’s established populations, from which Berrien County’s *A. americanum* likely originated, may have increased invasion success. Interestingly WDSP, where *A. americanum* appears to be establishing at a slower rate compared to GMSP, is closer to the Indiana populations. In fact, WDSP is ~48 km from the Indiana Dunes National Park and Indiana Dunes State Park, where *A. americanum* are abundant (L. Green, IDOH, pers. Comm., see below).

### Definition of established populations and importance of abiotic factors for establishment and invasion

The criteria for considering a tick species as being established in a given county is important to consider as we seek to understand their spread. Established tick populations as defined by the CDC involves either finding more than one life stage, or finding at least six ticks, in that county in a calendar year (Dennis et al. 1998, Springer et al. 2014, Nelder et al. 2019). Passive surveillance data also have been used to fulfill this criteria (Dennis et al. 1998, Springer et al. 2014). While community-submitted ticks can be an important source of data for informing risk, they can be unreliable for determining whether a tick population is established in a given area, if travel history is not taken into account and tick exposure may have occurred elsewhere in a tick endemic area. This may be a particularly important point for aggressive ticks such as *A. americanum* which can have multiple ticks feeding on a single host. A single dog traveling to an endemic area can easily bring back enough ticks to consider a county established. *Amblyomma americanum* submitted to the MDHHS for the last decade fulfill CDC criteria for some Michigan counties in some years, but the numbers vary widely from year to year, and no county is consistently represented (Figures 1-2). Additionally, active surveillance from those same years do not provide evidence for any established populations in those areas. More stringent criteria have been proposed by Nelder et al. 2019, specifically requiring that active surveillance is needed to confirm an established population. They further suggest that a population be confirmed for three consecutive years where at least six specimens from a given locale be collected, including at least one drag-sampled tick and at least one nymph. While this may seem like excessive criteria, it may be warranted, especially when establishment data are being used to train ecological niche models used to predict the northward spread of this vector in the coming years (e.g, Springer et al. 2015, Raghavan et al. 2019).

After lone star ticks are introduced into an area, both biotic and abiotic factors will affect establishment and invasion success. Given the generalist nature of *A. americanum,* and because one of their main hosts - the white-tailed deer - is very common and abundant in Michigan, invading *A. americanum* are not likely to be host-limited and instead may be more limited by mate-finding success, which will be overcome by increasing propagule pressure and off-host survivorship success. Abiotic factors, therefore, likely play a particularly critical role in population establishment along the northern edge of its distribution. These include foliage composition, temperature, humidity, day length and seasonality (Ludwig et al. 2016). Cold winters are known to limit the survivorship of overwintering ticks (Semtner and Hair 1976, Kaizer et al. 2015, Linske et al. 2019, Bacon et al. 2021), so it might not be purely coincidence that both in New England and the northern central U.S., populations of *A. americanum* are emerging initially along coastal habitats, which have more moderate winters relative to areas further inland. It is unknown how *A. americanum* may have been introduced into northwest Indiana, whether via terrestrial hosts or migrating birds, but there are records of low numbers of *A. americanum* collected via a combination of passive and active surveillance from the three northwest counties bordering Lake Michigan since the late 1990s and evidence of established populations since 2005 (L. Green, Indiana Department of Health, pers. Comm.). The Indiana Department of Health (IDOH) has been drag sampling *A. americanum* consistently for the last decade at the Indiana Dunes National Park and Indiana Dunes State Park, where there are robust populations (L. Green, IDOH pers. Comm.). Besides the moderating effect of Lake Michigan on winter conditions, the oak upland coastal dune forests along Lake Michigan provide sheltered areas with highly suitable microhabitat (i.e., thick leaf litter) that further insulate ticks from cold and desiccation (especially in the “valleys” on the leeward sides of dunes). *Ixodes scapularis* in Michigan’s Lower Peninsula likely invaded from northwestern Indiana and emerged faster northwards along Lake Michigan than it did farther inland (Hamer et al. 2010). Given *A. americanum*’s lower tolerance for cold winters, the invasion of this tick might follow a similar pattern. Additionally, if introduced, *A. americanum* may establish and emerge on the eastern side of the state along the Lake Erie and Lake Huron coastlines where the proximity to the lakes help create more moderate conditions facilitating winter survival.

Models have recently predicted that climate change resulting in longer summers and milder winters may increase the survival and therefore establishment probability of *A. americanum* farther north (Raghavan et al. 2019, Sagurova et al. 2019); correspondingly, these models predict that much of southern Michigan is already suitable for *A. americanum* tick establishment (Raghavan et al. 2019, Sagurova et al. 2019). It should be noted, however, that some ecological niche models (Springer et al. 2015, Raghavan et al. 2019) rely on passive surveillance data (Springer et al. 2014) that resulted in categorizing some Michigan counties as having established *A. americanum* populations, which as discussed previously, have not been confirmed through active surveillance (Walker et al. 1998). Sagurova et al. (2019), however, uses a mechanistic model based on regional climate and tick development rates, which does not rely on these data, and still predicts that *A. americanum* are likely to become established in Michigan in the immediate future.

### *Seasonal activity of* A. americanum *emerging in Michigan and implications for public health*

Frequent sampling at GMSP in 2020 allowed us insight into the phenologies of the three questing life stages of *A. americanum* in southwestern Michigan (Figure 3). Unfortunately, due to COVID-19 related restrictions, surveillance by MSU was unable to begin until 6/12/20 (although BCHD started surveillance 5/17/20, they did not detected any *A. americanum* at this site). The phenological trends, however, were consistent with that described in endemic areas further south and east (Sonenshine 1979, Schulze et al. 1986, Kollars et al. 2000, Carroll 2011, Gilliam et al. 2018). The seasonal patterns observed here are based on very low tick numbers. Assuming established *A. americanum* populations will continue to increase in abundance, continued monitoring encompassing early spring through fall will result in a greater confidence in observed patterns. Increased abundance also may result in broader activity periods than currently observed. For instance, perhaps more nymphs will be observed later in the summer, and some larvae may be detected in early summer as had been noted recently on Long Island and Massachusetts (Telford et al. 2019).

Although there are differences in phenologies between *A. americanum* and *I. scapularis*, the activity periods for all three life stages of *A. americanum* coincide with one of the major activity periods (i.e., larvae and adults) or overlap almost entirely (i.e., nymphs) with that of the respective life stages of *I. scapularis*. Coincident phenologies have practical implications for public health. First, efforts to survey for *I. scapularis*, which generally target the nymphal host-seeking period because of their preeminent epidemiological role, should also detect nymphal *A. americanum*, and to a lesser extent adults, thereby increasing the likelihood of detecting and characterizing emerging *A. americanum* in Michigan. Second and similarly, public health messaging that typically is targeted in spring to prevent and mitigate Lyme disease (e.g., promoting tick prevention and recognition of signs of a tick-borne disease) will also help reduce risk and negative impacts of *A. americanum*-associated diseases. Third, it is important for the public and for healthcare workers to recognize the different tick species and how they are associated with different diseases to prevent misdiagnosis and treatment of tick-borne diseases (e.g., over-diagnosis and treatment of Lyme disease and under-diagnosis and treatment of an *A. americanum*-associated disease) (Egizi et al. 2017). For example, post-tick antibiotic prophylaxis of Lyme disease (Lantos et al. 2021) can be provided for patients who detect a nymphal or adult *I. scapularis* that has fed for at least 36 hours and which had been acquired from a high Lyme disease incidence area, such as Berrien County (MDHHS 2021b). The coincident activity of *I. scapularis* and *A. americanum* nymphs, however, creates an additional challenge to determine the species of the small feeding nymphs, which thus may affect the decision to recommend post-tick antibiotic prophylaxis.

### Additional considerations for public health in emerging areas

*Amblyomma americanum* is of special concern in emerging areas as many of its associated diseases may not be familiar to medical professionals in the region. Pathogens of concern for human health that *A. americanum* are known to vector have been summarized in a recent review by Madison-Antenucci et al. 2020 and include *Ehrlichia chaffeensis* the causative agent of human monocytic ehrlichiosis (HME), *E. ewingii*, and the rare but highly fatal Heartland virus and Bourbon virus (Madison-Antenucci et al. 2020). For example, the numbers of *A. americanum* submitted to a passive surveillance program in central New Jersey have been increasing over the last decade and have overtaken that of *I. scapularis* in central New Jersey (Egizi et al. 2017, 2019). Yet relative to the estimated human encounter rates of *E. chaffeensis*-infected *A. americanum* to *Borrelia burgdorferi*-infected *I. scapularis*, there are significantly fewer expected reported cases of HME relative to Lyme disease, which may suggest an under-reporting or mis-diagnosis of HME cases (Egizi et al. 2017). Similarly, an increase in mild cases of spotted fever rickettsias (potentially caused by *Rickettsia amblyommatis*, which have been detected in a high proportion of *A. Americanum)* should be expected, but should not be mistaken for the much more severe Rocky Mountain spotted fever (Behravesh et al. 2016). One of the most concerning conditions associated with *A. americanum* is the recently discovered alpha-gal syndrome (AS). In *A. americanum* endemic areas, AS is the leading diagnosed cause of anaphylaxis (Pattanaik et al. 2018). One recent case study (Houchens et al. 2021) published in the *New England Journal of Medicine*, titled “Hunting for a Diagnosis,” describes a Michigan resident (with no travel history outside of the state) who arrived at an emergency room with severe anaphylaxis. After four days of hospitalization and multiple tests, the patient was diagnosed with AS. The details in this report demonstrate the potential severity of AS and the difficulty in diagnosing this unusual condition, especially in areas where *A. americanum* is newly emerging and awareness of AS may not be prevalent in the medical community (Houchens et al. 2021).

## Conclusion

Models have predicted that *A. americanum* would expand its range northward into Michigan (Molaei et al. 2019, Sagurova et al. 2019), but until 2019, active surveillance had not detected any established population. Here we report both an established *A. americanum* population at the northern extent of its range as well as the seasonal activity. Negative surveillance data in preceding years suggest that these findings most likely represent a nascent invasion rather than a long established but undetected population. Phenological patterns observed also provide insights into the public health risks posed by this tick over the course of the year and can help inform optimal time frames for future surveillance efforts. Due to limited resources and time, collected ticks have not yet been submitted for pathogen testing. Given similar host and habitat requirements, if the previous invasion of *I. scapularis* into Michigan is any indication (Hamer et al. 2010, Lantos et al. 2017, MDHHS 2020), we can expect to see *A. americanum* expanding its range in Michigan in the coming years. Educating public health officials and the medical community for both humans and companion animals about the diseases associated with this tick and how to differentiate them from other endemic tick-borne diseases is recommended.

## Acknowledgements

We thank S. Hamer, G.J. Hickling, and E.D. Walker for their contributions to beginning the long-term surveillance studies of invasive ticks in Michigan from which subsequent efforts have grown. Similarly, we thank E.D. Walker for providing additional insight into the data from Walker et al. 1998 as well as having coordinated and published that valuable work, which provides a baseline from which to compare subsequent distributions of ticks over time. For assistance in the field and lab, we thank S. Altus, I. Arsnoe, M. Bammer, A. Dunivant, K. Fake*, S. Froehlich, M. Gleason*, G. Grzesiak, S. Haupt*, D. Houvener, S. Kim, J. Kryda*, A. Larson, A. Luchenbill, M. Orbain, G. Pang, J.Pastori, M. Regalado, B. Reith, M. Rice, M. (Rosen) Clayson, J. Schroeder, M. Soja, N. Spala, J. Stych, A. Talbott, M. Volk, H. Waters, K. Wickens, B. Wilson*, A. Yackley, G. Yarandi, S. Zohr, and many other assistants who contributed to earlier, published work. We thank our agency partners who variously started and continue to maintain the Michigan Department of Health and Human Services (MDHHS) community tick submission program; developed, funded, and facilitated active invasive vector surveillance efforts conducted by local health departments; and provided suggestions for MSU active surveillance efforts based on public health risks. They include R. Eisen, M. Poplar, R. Reik, J. Sidge, K. Signs, and M.G. Stobierski. We furthermore thank L. Green from the Indiana Department of Health (IDOH) for providing historical knowledge and guidance on sampling for *A. americanum* in northern Indiana. Permission to sample properties was graciously provided by the Michigan Department of Natural Resource, National Park Service, many county parks offices, nature centers, land conservancies, universities, and other public and private entities. We are grateful for many sources of funding supporting the work presented here. First and foremost, the majority of the active surveillance efforts for invasive ticks in the Tsao lab has been generously supported by the Michigan Lyme Disease Association (MLDA), facilitated by the L. Lobes. Drag surveillance by the Tsao lab in 2019 was largely funded by Contract E20193274-00 from MDHHS. Funding for the companion animal surveillance study was provided by the College of Veterinary Medicine Endowed Companion Animal Fund RT082792, the MLDA, and greatly facilitated by L. Penman and Boehringer Ingelheim Animal Health. Drag surveillance by the Berrien County Health Department was funded in part by the MDHHS, and the MDHHS community tick submission program is supported with funds from the Epidemiology and Laboratory Capacity (ELC) for Prevention and Control of Emerging Infectious Diseases Cooperative Agreement from the Centers for Disease Control and Prevention, award number CDC-RFA-CK19-1904. We thank the Michigan State University (MSU) Comparative Medicine and Integrative Biology program for fellowship funding for PF, MP, and VK. MP, VK, and JT also have been funded by the CDC Midwest Center of Excellence in Vector-Borne Diseases U50723K866 under a subcontract to Michigan State University from the University of Wisconsin (L. Bartholomay, S. Paskewitz, principal investigators). This publication was supported by Cooperative Agreement #U01 CK000505, funded by the Centers for Disease Control and Prevention including funding for SN, LQ, and several other research fellows listed above with asterisks (*) who contributed to surveillance efforts. Its contents are solely the responsibility of the authors and do not necessarily represent the official views of the Centers of Disease Control and Prevention or the Department of Health and Human Services.

